# Solriamfetol enhances alertness and cognitive performance in mice

**DOI:** 10.1101/2025.01.02.631072

**Authors:** Meriem Haddar, Stamatina Tzanoulinou, Li Yuan Chen, Mehdi Tafti, Anne Vassalli

## Abstract

Solriamfetol [(R)-2-amino-3-phenylpropylcarbamate hydrochloride], a phenylalanine derivative initially developed as potential antidepressant, was shown by our group in 2009 to have potent, dose-dependent wake-promoting activity in mice. Solriamfetol (Sunosi^®^) is used since 2019 to counteract excessive daytime sleepiness (EDS) in patients with narcolepsy and obstructive sleep apnea (OSA). It has several advantages over other stimulants, notably that it is not associated with strong psychomotor activity, does not induce behavioral stereotypies and anxiety-related behaviors, in contrast to amphetamines and modafinil. Its mode-of-action remains incompletely solved. It was reported to act as dual dopamine-and-noradrenaline-reuptake-inhibitor (DNRI), and more recently, to have TAAR1 agonist activity. In our early mouse study, we showed that, at 150 mg/kg, Solriamfetol induces a state of wakefulness featuring a dramatic upregulation of EEG gamma activity, a cognitive biomarker, and the expression of genes implicated in neural plasticity, learning and memory. Besides being prescribed to treat EDS, central nervous system stimulants are commonly used as ‘smart drugs’ to enhance cognition in normal individuals. Therefore, based on Solriamfetol’s ability to potently induce wakefulness, EEG and molecular markers of learning and memory, we aimed to determine whether it could improve the cognitive performance of wild-type mice. Because at doses of 50-150 mg/kg, Solriamfetol induced an alert waking state associated with low mobility, thus precluding behavioral testing, we used lower doses (1-3 mg/kg) to assess cognition in a battery of tests evaluating short- or long-term memory and spatial navigation. We found that compared to saline, Solriamfetol 3 mg/kg consistently improves sustained attention for a novel object, as well as spatial memory. Next, to determine the brain activity correlates of enhanced cognition, we performed EEG/EMG recording while the mice performed the novel object recognition (NOR) task. Power spectral density (PSD) analysis revealed that 3 mg/kg Solriamfetol reduced EEG delta (an index of sleepiness) during exposure to a novel context, and enhanced EEG fast-gamma (an index of mental concentration) during execution of the NOR task. Taken together, our data demonstrate that low-dose Solriamfetol improves memory and attentional performance in wild-type mice.

## Introduction

Excessive daytime sleepiness (EDS) is a debilitating and highly prevalent condition with profound morbidity. It is present in a wide range of neurological diseases, including central hypersomnia, such as narcolepsy, idiopathic hypersomnia, and Kleine-Levin syndrome, circadian rhythm sleep-wake disorders, breathing-related sleep disorders, such as obstructive sleep apnea (OSA), and is also common in other neurological disorders, including Parkinson’s disease, myotonic dystrophy and multiple sclerosis, and in psychiatric disorders, such as major depressive disorder and stress-associated disorders.

The causes of EDS are varied, and therapy must be tailored accordingly. Central nervous stimulants (CNS) increase alertness, attention, physical activity, and energy levels through diverse mechanisms believed to alter the levels of critical brain neurotransmitters. The most commonly used, caffeine, potently antagonizes adenosine signaling, which causes drowsiness. Thus, by reducing adenosine activity, caffeine increases energy levels, but its use typically triggers a powerful subsequent sleepiness rebound. The first medically discovered stimulant was amphetamine^1^. With its derivatives such as methamphetamine, amphetamines increase catecholamine (dopamine and noradrenaline) release and inhibit their reuptake. These compounds are reported to be highly addictive, and they cause acute and potentially long-lasting side effects such as hypertension^2,3^.

For decades, modafinil has been the most widely used wake-promoting drug to alleviate EDS, especially in narcolepsy. Modafinil is believed to act through the dopaminergic system although its precise mode of action is not fully solved^4-6,7^. Modafinil can however have side effects, and a significant number of narcolepsy patients are modafinil-unresponsive^8,9^. Although most stimulants have a major dopaminergic-based mode of action, alternative stimulants are being developed, that affect the levels of norepinephrine, histamine, serotonine, or trace amino acids (TAAs).

First investigated as potential antidepressant, the phenylalanine derivative Solriamfetol, was later abandoned for this indication. Among the most significant effects found in human studies was insomnia^10^. In 2009, our group demonstrated that Solriamfetol administered at 150 mg/kg in early light (i.e., sleeping) phase acted as a potent wake-promoting agent in 3 different strains of mice^11^. And indeed, since 2019, Solriamfetol (Sunosi®) is approved to counteract EDS in narcolepsy and obstructive sleep apnea (OSA) patients.

Mechanistically, Solriamfetol is thought to act as a dual dopamine and noradrenalin reuptake inhibitor (DNRI)^12^. Recently, Solriamfetol was reported to additionally have TAAR1 agonist *in vitro* binding activity, unlike modafinil and bupropion, but in common with amphetamine and methamphetamine, as well as 5HT1A receptor *in vitro* binding activity^13,14,15^.

Besides demonstrating wake-promotion, our 2009 murine study additionally demonstrated that Solriamfetol has unique properties relative to d-amphetamine and modafinil on the quality of the induced wakefulness, that featured a delayed increase in EEG theta power, as well as a dramatic enrichment in slow- (35-60 Hz) and fast- (60-80 Hz) EEG gamma activity during 1-4 h following drug administration^11^. Both EEG theta and fast-gamma oscillations are correlates of mental concentration, believed to play instrumental functions in cognition^16,17^. Furthermore, Solriamfetol induced hippocampal expression of *c-Fos*, as well as plasticity-related genes, including *Crebbp, Gsk3b, Ncan, Ntrk2, Ptprd* and *Vldlr* that show upregulation in assays of long-term-potentiation and other hallmarks of learning and memory^11^. Altogether, upregulation of EEG gamma and induction of neuronal plasticity-related and immediate early genes, raised the question of whether Solriamfetol may have cognitive enhancing properties. Besides being prescribed to treat EDS, CNS stimulants are also used as ‘smart drugs’, in particular modafinil to enhance cognition in normal individuals.

Solriamfetol reduces anxiety-like behavior in mice, and restores the deficit in object preference induced by intermittent hypoxia^18^. Similar results were found with mice that underwent sleep fragmentation^19^. However, in these two earlier studies, mice were under intermittent hypoxia or sleep fragmentation, and Solriamfetol was administered at high doses at dark onset, hours before the behavioral experiments that were performed during the light period.

The present study thus aimed at determining whether Solriamfetol could improve cognitive performance in wild-type mice. Because at 150 mg/kg, 100, or 50 mg/kg, Solriamfetol was found to induce a state of wakefulness associated with low mobility lasting for ∼1-2 hours post injection, and therefore precluded behavioral testing, we used lower doses (1 or 3 mg/kg) to assess cognition in a battery of tests that evaluate short- or longer-term memory, as well as spatial navigation. We found that Solriamfetol at 3 mg/kg consistently improves sustained attention for a novel object, as well as spatial memory, compared to saline injections.

## Materials and Methods

### Animals

Thirty-six C57BL/6J male mice were obtained from Charles River, France, and housed in standard conditions (12 h/12 h light/dark cycle), with light-onset at 07:00 AM, defined as Zeitgeber zero (ZT0), with *ad libitum* access to water and standard rodent chow. All animal procedures followed Swiss federal laws and were approved by the State of Vaud Veterinary Office. Care was always taken to optimize wellbeing and minimize discomfort and stress.

### Animal treatment and drug administration

When mice reached 11-12 weeks of age, they were first habituated to handling and intraperitoneal (i.p.) saline injection for 3 days, and then randomly ascribed to one of 3 groups to undergo a 12-day pipeline of behavioral tests after daily saline (n=12 mice), Solriamfetol 1 mg/kg (n=12 mice), or Solriamfetol 3 mg/kg (n=12 mice) administration (Fig. 1A).

**Figure 1.**
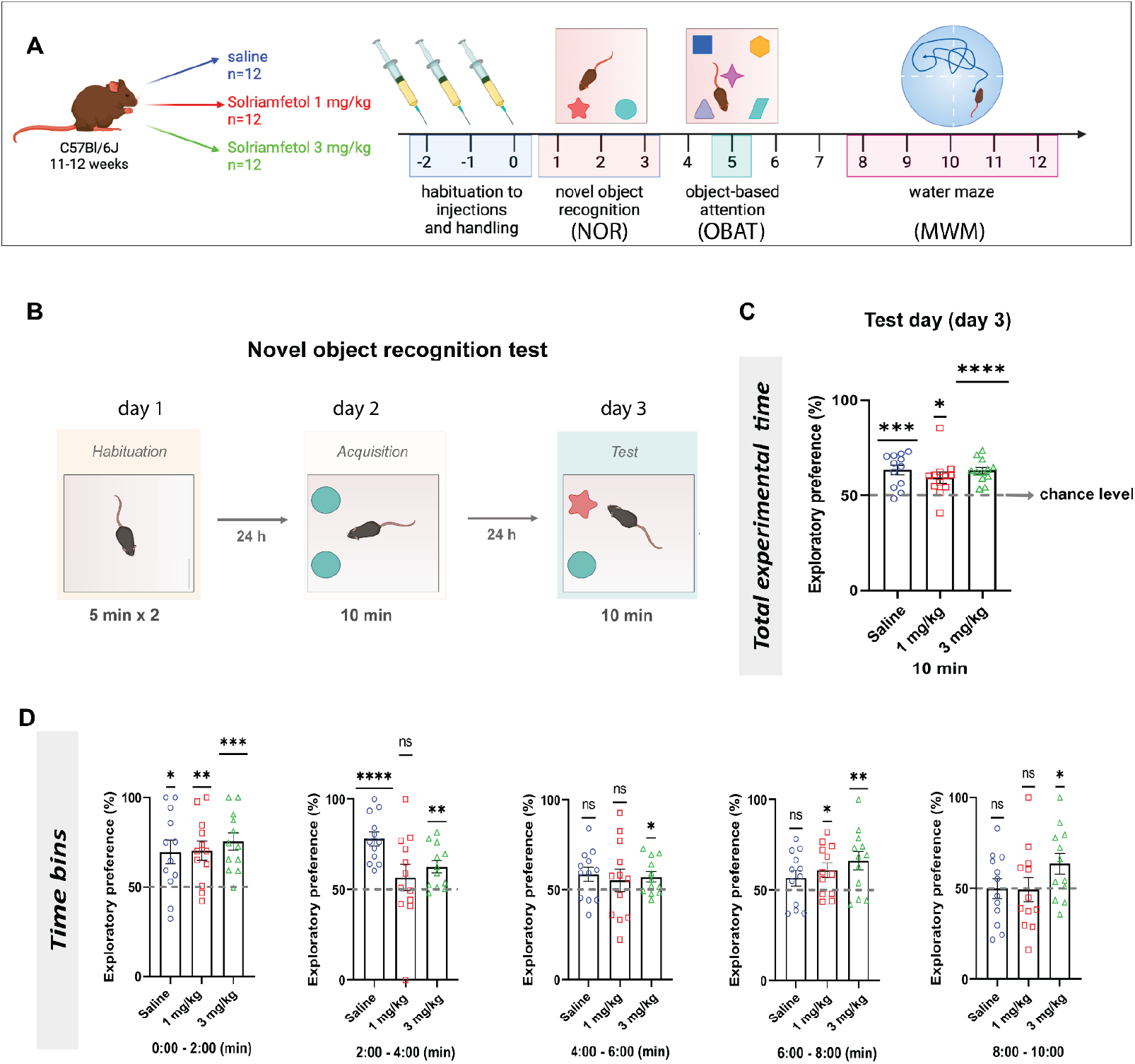
Effects of Solriamfetol on performance of C57BL/6J wild-type mice in the novel object recognition (NOR) test. A. Schematics depicting the behavioral pipeline used to determine Solriamfetol effects on the cognitive performance of wild-type mice. Drug or saline was administered intraperitoneally every day ∼30 min prior to the start of the behavioral sessions. **B**. Schematic representation of the NOR test. **C**. Exploratory preference for the novel object during the 10 min test on day 3. The dashed line represents chance level (50%). One sample t-test compared to chance level, \**P*<0.5, \*\*\**P*<0.001, \*\*\*\**P*<0.0001. n=12/group. Solriamfetol 1 mg/kg or Solriamfetol 3 mg/kg groups vs Saline group: one-way ANOVA followed by Dunnett’s multiple comparisons test: F(2, 33) = 0.819, *P*>0.05. **D**. Dynamics of the exploratory preference for the novel object dissected in 2-min time-bins. As the 10 min NOR test progresses, the Saline and Solriamfetol 1 mg/kg groups tend to lose preference for the novel object, while Solriamfetol 3 mg/kg is the only group to preserve preference for the novel salient object throughout all time bins. One sample t-test compared to chance level, *\*P*<0.5, \*\**P*<0.01, \*\*\**P*<0.001, \*\*\*\**P*<0.0001. ns, non-significant (*P* > 0.05). n=12 mice/group. Data are expressed as mean ± SEM.

To replicate a chronic daily medication regimen in humans, Solriamfetol or saline were injected daily across the 12-day experimental pipeline (see Fig. 1A), including in the days without behavioral tasks. Solriamfetol was provided by JAZZ Pharmaceuticals (Dublin, Ireland), stored at room temperature, and it was freshly prepared daily by dissolution in saline (0.09% NaCl). The dose was adjusted to the mouse body weight and administered i.p. (5 μl of solution per body gram). Saline or Solriamfetol were administered 30 min before the start of each behavioral task, or at the same time of the day on non-experimental days. Tests were performed in the Light Phase (generally between 9 AM and 2 PM, i.e., ZT2-7), to allow mice to view the visual cues used for the Morris Water Maze (MWM) test.

Animals used for EEG/EMG recordings were similarly treated, except that the injections were performed at ZT12 and the novel object recognition test (NOR) was performed shortly after dark-onset (ZT12.5-13), i.e., in early active phase.

### Novel object recognition (NOR) test

The NOR test was used to investigate object recognition memory and consisted of 3 experimental days. On the 1^st^ day the mice were habituated to the test arena (44 cm X 44 cm) for 2 x 5-min trials with an inter-trial-interval (ITI) of 30 min (habituation phase). On the 2^nd^ day, each animal was placed in the same arena, now containing 2 identical objects (10 cm away from the walls) and allowed to explore for 10 min (acquisition or learning phase). After a 24 h delay, the animal was placed in the same arena with one of the previously explored objects replaced by a novel one and allowed to explore for 10 min (test phase). The exploratory preference (%) was calculated as the percentage of time spent exploring the novel object (N) divided by the sum of exploration time of the novel and the familiar object (F): exploratory preference (%) = (N/(F+N)) x 100. Object exploration time, defined as time during which the tip of the animal head was within ∼2 cm of the object, was manually scored.

To conduct the NOR test in mice instrumented for EEG/EMG recordings, a different cage setup was used (37.3 cm X 23.4 cm) with an added plexiglass floor.

### Object-based attention test (OBAT)

OBAT consisted of 2 phases carried out in one single day: an acquisition phase and a retention phase. During the acquisition phase, animals are placed during 4 min in an arena containing 5 objects separately positioned in the central zone of the arena. The mouse is then placed in a waiting cage, while 4 out of the 5 objects are removed and replaced by one novel object. In the retention phase, the mouse is placed again in the arena and allowed to explore the 2 objects (one familiar and one novel) during 5 min. The exploratory preference is expressed as above as exploratory preference = (N/(F+N)) x 100, where N and F are the time spent during the retention phase within ∼2 cm of the novel (N) and familiar (F) object, respectively^20^.

### Morris Water Maze (MWM) test

The water maze test was used to assess spatial learning and memory and consisted of 5 experimental days. The water maze consists of a circular pool (150 cm diameter), filled with opaque water. The temperature is maintained at 25°C ± 1°C during the experiment. A platform is submerged under the water surface, and therefore not visible. However, once the mouse finds its location, it allows the animal to stop swimming and rest. As mice are averse to swimming in water, finding the platform is a strong motivation. The pool is surrounded by visual cues that the animal can use to learn to navigate as the training progresses. On the first day of the MWM experiment, mice were accustomed to the pool for two sessions each (2 x 60 s). During this habituation, the platform was placed in the middle of the pool without any cues being used. The learning sessions were conducted for 4 consecutive days, during which each animal was subjected to 4 x 60-s trials per day (days 1, 2, 3, 4) with an interval of 15-20 min between 2 consecutive trials. If a mouse fails to find the platform within 60 s, it is softly guided and placed on it. Each mouse must stay on the platform for 20 s, after which it is returned to a holding cage until the next trial. The probe trial is performed 24 h after the last training day (day 5). During the probe, the platform is removed, and the mouse is allowed to swim freely for 60 s. The percentage of time spent swimming in the quadrant that contained the platform during training (target quadrant) was measured and analyzed, offering an index of spatial memory.

### Electroencephalography/Electromyography (EEG/EMG) recording

On the day of surgery, mice were anesthetized using 5% isoflurane/100% O_2_ gas mixture and then fixed on a stereotaxic frame. Isoflurane was then gradually reduced to 2.5%, which allowed to maintain general anesthesia throughout the surgery. Immediately after fixing the mouse head in the stereotaxic frame, Carprofen (Rimadyl, 5 mg/kg; Zoetis, Delemont, Switzerland) was administered subcutaneously. The skull was then exposed and 4 gold-plated screws (J. I. Morris, Oxford, MA, USA), soldered beforehand with gold wire, were screwed into the skull. 2 screws were used as electrodes and signal sensors (anteroposterior [AP] and mediolateral [ML] distance with reference to bregma): the Frontal (AP: +1.0 mm, ML: +1.0 mm) and Parietal (AP: -3.2 mm, ML: +1.0 mm) electrodes. Another screw served as ground electrode (AP: -1.0 mm, ML: -2.0 mm). The 4^th^ screw was implanted on top of the cerebellum area and used as reference (AP: -5.2 mm, ML: -2.0 mm). The screws were attached to the skulls with a self-adhesive resin cement (3M™ RelyX™ Unicem 2 Automix Self-Adhesive Resin Cement; DentoNet AG, Zurich, Switzerland). Afterwards, gold wires of these screws were threaded through and soldered to the 3 holes of a Pinnacle EEG/EMG headmount (8201-SS, Pinnacle Technology Inc., Lawrence, KS, USA). 2 stainless steel wires were inserted in the neck muscles to serve as EMG electrodes and attached to the headmount. Finally, the implant was sealed together using an autopolymerizing prosthetic resin (Paladur; Kaladent, Lausanne, Switzerland). Mice were postoperatively individually housed, with paracetamol (Dafalgan, 1 mg/ml; UPSA, Zug, Switzerland) dissolved in drinking water for 1 week. After 5 days of post-operative recovery, mice were habituated to the recording cables of the Pinnacle acquisition system for 5 further days. After habituation, EEG/EMG data collection was initiated using Sirenia Acquisition (Pinnacle Technology), with EEG and EMG signals high-pass filtered above 0.7 and 10 Hz, respectively, and sampled at 2,000 Hz. Recordings started with 2 baseline days, followed by a day with saline injection (i.p.) at ZT12, then by 6 consecutive days with daily saline or Solriamfetol (3 mg/kg) injection at ZT12, for the Saline or Solriamfetol groups, respectively. During the last 3 days, the NOR test was performed as described above, starting 30 min after injection.

### Vigilance state scoring and analysis

EEG/EMG recordings were divided into 4-s epochs and vigilance states were visually scored as wakefulness, non-rapid eye movement (NREM) sleep, or rapid eye movement (REM) sleep, using previously described criteria^21,22^ and the SleepSign software (KISSEI COMTEC CO, LTD, Nagano, Japan). State scoring events were exported as Excel files, and EEG signals linked and analyzed using Brainstorm^23^ and custom MATLAB scripts. EEG signals were downsampled to 400 Hz. The 50 Hz powerline artefact was removed by notch filtering EEG signals between 49 and 51 Hz. EEG signals were subjected to Fourier transform (non-overlapping 4-s windows, 0.25 Hz frequency-bin resolution with Hamming window) to determine EEG power spectral density (PSD) of wakefulness across 0.25–200 Hz. Adapting from Hasan et al^11^, EEG spectra during Saline or Solriamfetol injection days were expressed as percentage of baseline, which was calculated for each mouse by averaging the EEG PSD in each frequency bin between ZT12-16 across both baseline days. The PSD during behavior was extracted to compare the Saline and Solriamfetol groups. The PSD was averaged by frequency windows: delta (1-4 Hz), inter-delta/theta (4-7 Hz), theta (7.5-11.5 Hz), beta2 (23-30 Hz), slow-gamma (32-45 Hz) and fast-gamma (55-85 Hz).

### Statistical analysis

Statistical analyses were performed using GraphPad Prism 9.1.2 (GraphPad Software, Boston, Massachusetts, USA). Two-tailed T-Test, one sample T-Test against chance level, one-way ANOVA followed by Dunnett’s multiple comparisons test, two-way ANOVA followed by Dunnett’s multiple comparison tests, repeated measures two-way ANOVA followed by Dunnett’s multiple comparisons test or mixed-design ANOVA followed by Greenhouse-Geisser correction and Šídák’s multiple comparison were performed. These analyses were performed based on the tests and their design (see Figure legends).

## Results

We previously reported that Solriamfetol administered in early resting phase, i.e., at light onset or ZT0, potently induces wakefulness in a dose-dependent manner. At 150 mg/kg, it delays sleep onset for 4-5 h (Hasan et al., 2009). This potent wake-promoting effect is however accompanied by an unusual effect on locomotion for the first ∼1 h. During this interval, Solriamfetol 150 mg/kg induces a state of wakefulness, when the mice are alert and attentive, open-eyed and responding to stimuli, but manifesting very limited spontaneous locomotor activity. This is in strike contrast with other CNS stimulants such as d-amphetamine, which induce hyperactivity with stereotypic movements such as licking and climbing the sides of the cage. At the reduced doses of 100, or 50 mg/kg, we observed that Solriamfetol still had negative effects on locomotion and exploration. As rodent cognitive tests rely on locomotor activity, reduced locomotion precludes assessment of cognitive performance. We found that lower doses of Solriamfetol (1 or 3 mg/kg) maintained movement, and we therefore asked whether these lower doses would enhance performance in learning and memory tasks. 36 C57BL/6J mice were thus divided into 3 groups, aimed at receiving saline (Saline group, n=12), Solriamfetol 1 mg/kg (n=12), or Solriamfetol 3 mg/kg (n=12. After 3 days of habituation to handling and i.p. injection of saline, the 3 groups of mice were led through a sequential battery of behavioral tests, evaluating different aspects of cognition during a total of 12 days (Fig. 1A).

### Solriamfetol enhances cognitive performance

Novel Object Recognition (NOR) test was first used to assess long term (24 h) memory for inanimate objects. Mice being innately highly curious for novelty, tend to explore novel objects that are neither-aversive, nor-appetitive, preferably to familiar ones. All 3 groups showed preference for the novel object over the familiar one (exploratory preference significantly above chance level of 50% in all 3 groups) when the total test duration of 10 min was considered (Figure 1C). However, when dissecting the total exploratory time into 2-min time-bins and comparing the novel object preference to chance level (50 %), the Solriamfetol 3 mg/kg group was the only one to maintain preference for the novel object relative to the familiar one throughout the five 2-min time bins of the NOR test (Figure 1D). As the 10 min of the test progress, the Saline and Solriamfetol 1 mg/kg groups tend to lose preference for the novel object (see last 2 min), while the Solriamfetol 3 mg/kg group preserves preference for the novel salient object throughout the 10 min, suggesting an effect of Solriamfetol on sustained attention for saliency.

Because learning performance of WT C57BL/6J mice may already be close to optimal in this test (i.e., ceiling effect), thus preventing us from detecting a robust drug-induced enhancement, we chose as the 2 next behavioral assays, more cognitively demanding tasks, namely object-based attention, a test requiring exploration of a higher number of objects in a shorter time, and the Morris water maze task, where mice undergo shorter training sessions.

We thus next performed the object-based attention test (OBAT), requiring a more elaborate set of attentional skills. Here, the mice are exposed to 5 different objects for 4 min (acquisition phase), then placed shortly in a waiting cage, while 4 of the 5 objects are removed and one novel object is added. The mouse is then immediately placed back in the arena for 5 min (test phase) (Figure 2A). We found that none of the 3 groups were able to demonstrate novel object preference over chance level during these 5 min, confirming the highest difficulty of the OBAT vs the NOR test (Figure 2B, left panel). Nevertheless, mice treated with Solriamfetol at 1 mg/kg and 3 mg/kg maintained a significantly greater distance between the tip of their nose and the familiar object than did saline-treated mice (Figure 2B, right panel), suggesting that the familiar object had less potential interest than the novel object. Moreover, when the 5 min were analyzed in 1 min-time-bins, the Solriamfetol 1 mg/kg and 3 mg/kg groups showed a tendency towards greater exploratory preference for the novel vs familiar object during the late phase (the last min) of the task compared to the saline group (*P*=0.054 and *P*= 0.057 Figure 2C).

**Figure 2.**
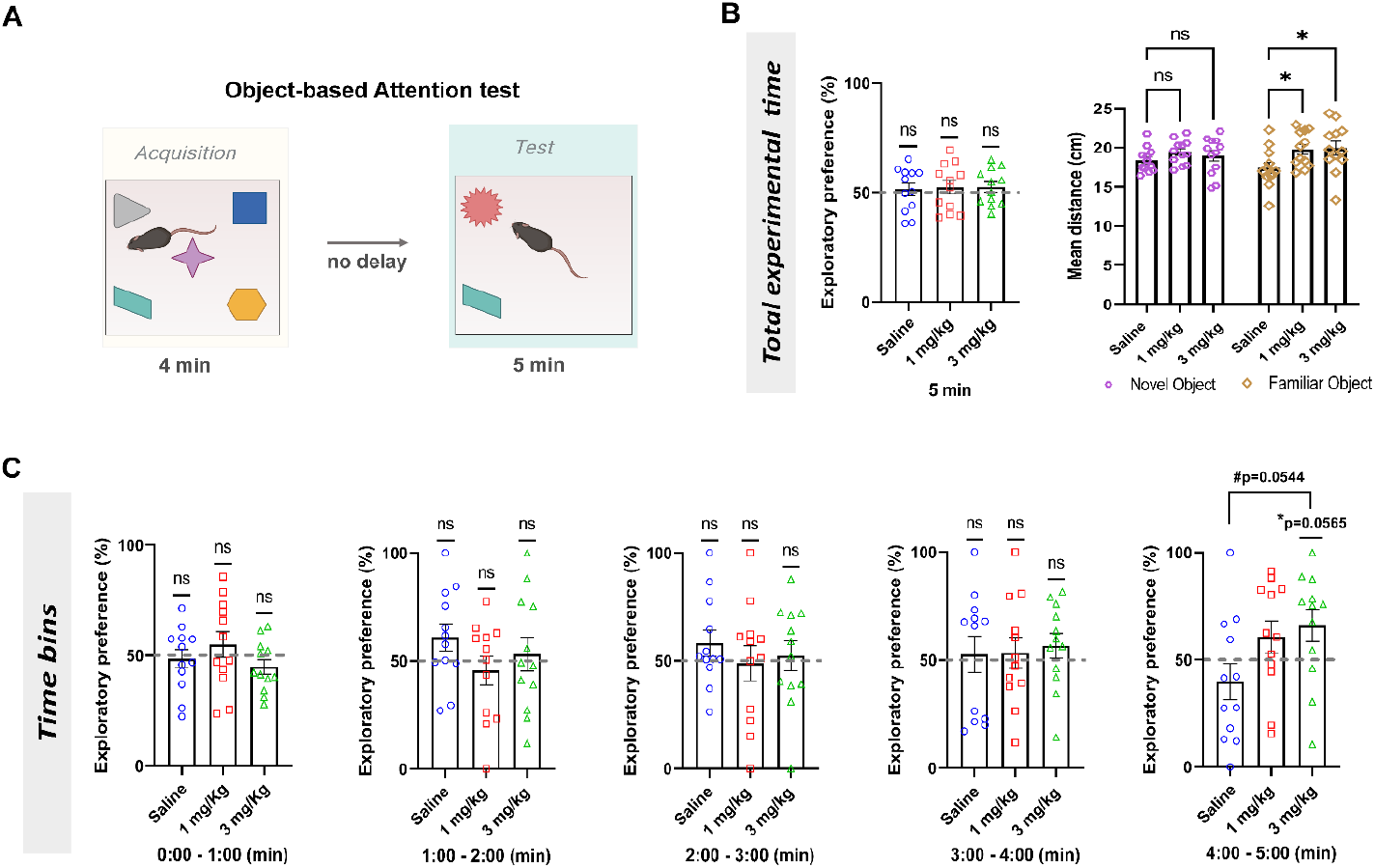
Effects of Solriamfetol on performance in the object-based attention test (OBAT). A. Experiment schematic. Drug or saline was administered intraperitoneally ∼30 min prior to the start of the behavioral test. **B**. Exploratory preference for the novel object (left panel) and mean distance to both objects during the 5 min test duration. Left panel: one sample t test, ns, non-significant effect compared to chance level (50 %). Solriamfetol 1 mg/kg and Solriamfetol 3 mg/kg groups vs Saline group: one-way ANOVA followed by Dunnett’s multiple comparisons test: F(2, 33) = 0.046, *P*>0.05. Right panel: 2-way ANOVA followed by Dunnett’s multiple comparison tests: drug effect, *P*=0.020, F(2, 66) = 4.13, no interaction effect. \**P*<0.05, compared to Saline group. **C**. Dynamics of the exploratory preference for the novel object dissected into 1-min time-bins. Repeated measure two-way ANOVA followed by Dunnett’s multiple comparison. No main effect. *P*=0.054 compared to Saline group. One sample t test. *P*=0.057, ns, not significant against chance level (50%). n=12 mice/group. Data are expressed as mean ± SEM.

We next conducted the MWM test, which assesses learning and memory during spatial navigation. The test spans 5 consecutive days, with pool habituation on the 1^st^ day, then 3 days for learning the platform location (with 4 daily sessions), and 1 final probe day in absence of platform (Figure 3A). We found no differences between the 3 mouse groups in total distance traveled by swimming during the 12 learning sessions (Figure 3B). Nevertheless, counting the number of mice that swam across the platform zone during the probe day revealed a dose-dependent effect: the number of mice swimming across the target zone for the Saline, Solriamfetol 1 mg/kg and 3 mg/kg groups were, respectively, 4 (33.33%), 6 (50%) and 8 (66.66%), out of 12 mice per group (Figure 3C). Although, the 3 groups did not show a significant preference for the target quadrant zone during the total probe time (1 min, Figure 3D), the Solriamfetol 3 mg/kg was the only group showing a trend for preference above the 25% chance level during the late phase of the probe trial when time was divided into 30 s bins (Figure 3E). Next, we assessed the distance traveled in the target quadrant and found that, during the last 30 s of the probe, the Solriamfetol 3 mg/kg group showed a significantly higher distance swam in the target quadrant than the 2 other groups (Figure 3F). To rule out that this was not due to an increase in locomotor activity of the 3 mg/kg group, we calculated the percentage of the total distance traveled across the target quadrant, and found that the 3 mg/kg group does show a tendency for increased % time in target quadrant during the last 30 s of the probe trial compared to the Saline group (Figure 3G, *P*=0.054). Chi-square analysis revealed that the fraction of mice exploring the target quadrant above chance level (25%) was higher in the Solriamfetol 3 mg/kg group than in the other 2 groups (9/12 vs. 5/12 and 3/12; Figure 3H, chi-square, *P*=0.044).

**Figure 3.**
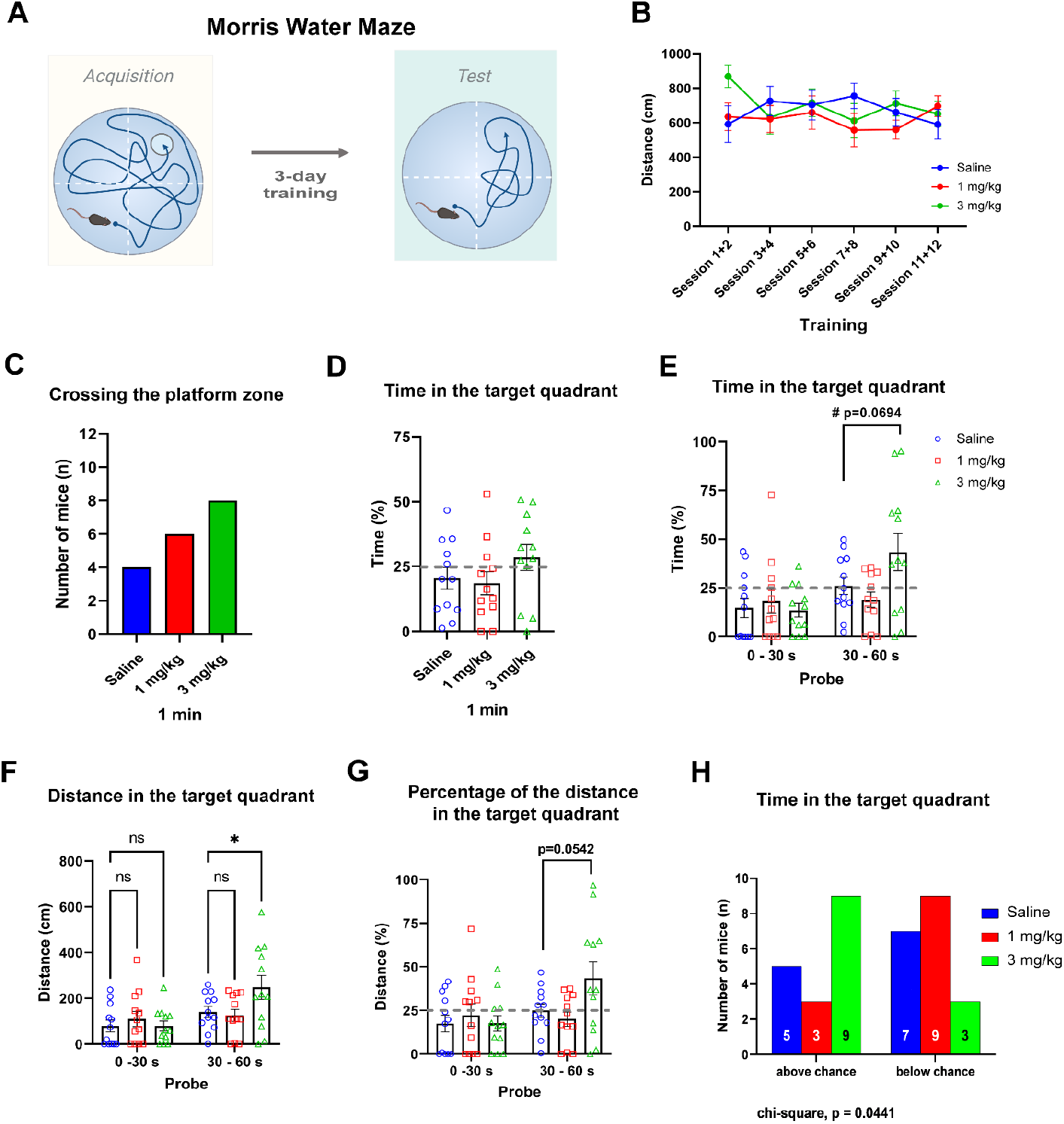
Effects of Solriamfetol on performance in the Morris water maze (MWM) test. A. Experiment schematic. Drug or saline was administered intraperitoneally on each of the 5 days ∼30 min prior to the start of the behavioral sessions. **B**. Total distance traveled in the maze during training. Repeated measures 2-way ANOVA followed by Dunnett’s multiple comparisons test, no main effect, not significant. **C**. Number of mice swimming across the virtual zone above the platform location during the probe trial. **D**. Percent of time spent in the target quadrant during the total duration of the probe trial. One-way ANOVA, not significant compared to Saline group. **E**. Time in the target quadrant during the probe trial divided in two time-bins. Repeated measures two-way ANOVA followed by Dunnett’s multiple comparisons test. TreatmentXTime interaction effect: F(2, 33) = 4.239, *P*<0.05. Time effect: F(1, 33) = 11.51, *P*<0.01. Treatment effect: F(2, 33) = 1.342, *P*>0.05. Dunnett’s Post-hoc comparison: *P*=0.069 Solriamfetol 3 mg/kg compared to Saline group. **F**. Distance traveled in the target platform during the probe trial divided by two time-bins. Repeated measures two-way ANOVA followed by Dunnett’s multiple comparisons test. Treatment x Time effect: F(2, 33) = 3.835, *P*<0.05. Time effect: F(1, 33) = 12.67, *P*<0.01. Treatment effect: F(2, 33) = 1.339, *P*>0.05. Dunnett’s Post-hoc comparison: \**P*<0.05, ns non-significant, compared to Saline group. **G**. Percentage of the distance traveled in the target platform during the probe trial divided by 2 time-bins. Repeated measures two-way ANOVA followed by Dunnett’s multiple comparisons test. DrugXTime interaction effect: F(2, 33) = 3.720, *P*<0.05. Time effect: F(1, 33) = 5.852, *P*<0.05. Treatment effect: F(2, 33) = 1.391, *P*>0.05. Dunnett’s Post-hoc comparison: *P*=0.054 Solriamfetol 3 mg/kg compared to Saline group. **H**. Number of mice exploring the target quadrant during the probe trial above (left) and below (right) chance level (i.e. 25%). Chi-square test. *P*=0.0441. n=12 mice/group. Data are expressed as mean ± SEM.

### Solriamfetol enhances EEG markers of alertness and concentration during the NOR test

To next determine whether enhanced cognitive performance is associated with changes in brain cortical activity, we performed the 3-day NOR test in mice instrumented for EEG/EMG recording. Unlike the NOR test above performed during the day (Figure 1), the drug is here administered at dark-onset (ZT12) on each of the 3 days, i.e., in the early active phase of mice, to emulate the human situation.

30 min after saline or Solriamfetol 3 mg/kg injection, mice underwent behavioral testing. EEG power spectral density (PSD) of the waking state was assessed 10 min before, during, and 10 min after each test phase (in 1 min time-bins, Figure 4).

**Figure 4.**
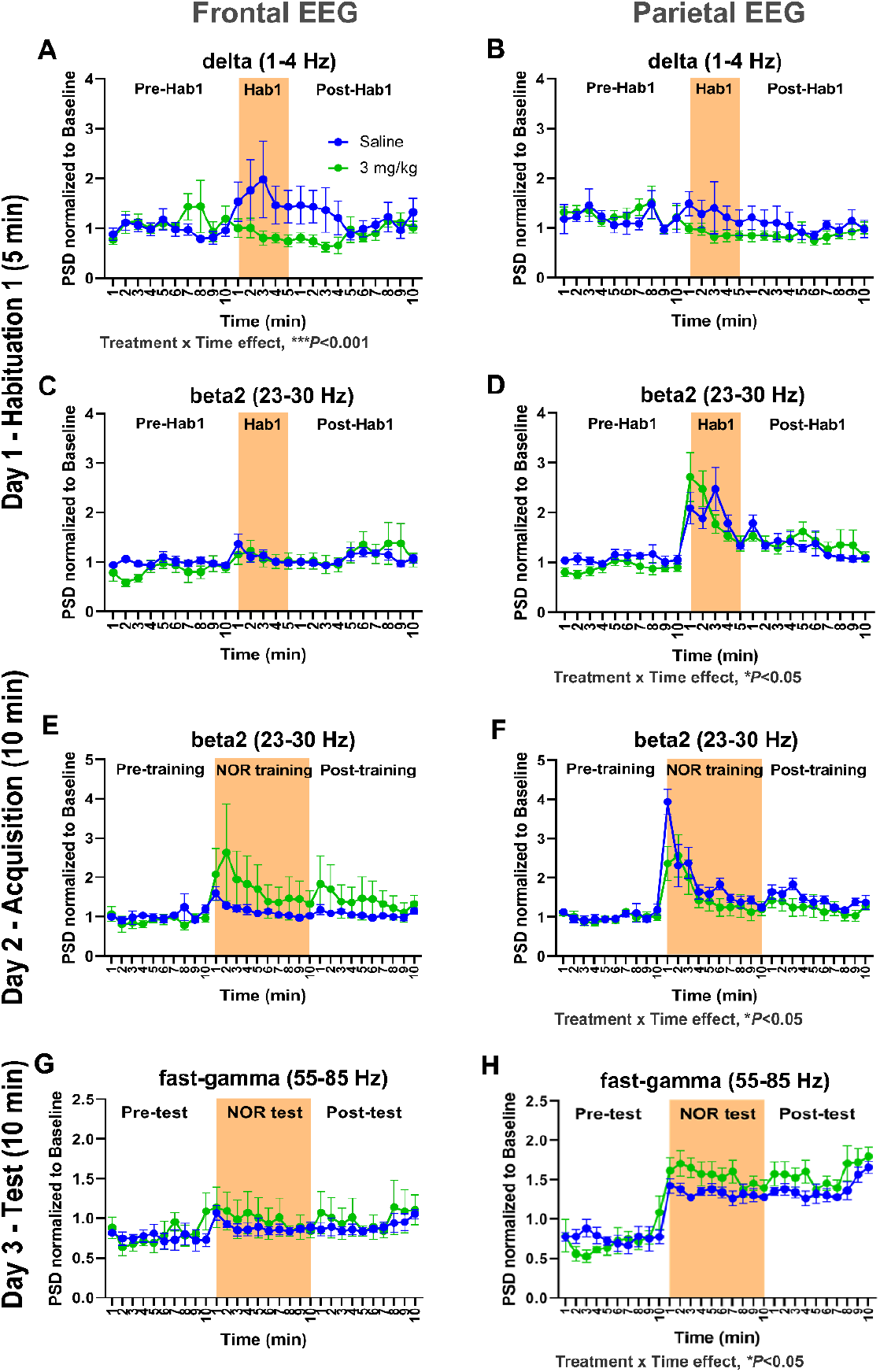
Effects of Solriamfetol (3 mg/kg) on the waking EEG of C57BL/6J mice during the NOR test. Saline or Solriamfetol 3 mg/kg were administered at dark-onset (ZT12) on each day of the 3-day NOR test, and the behavioral sessions started 30 min later. **A-B** depict the frontal (A) and parietal (B) delta power before, during and after the first 5-min habituation phase to the test arena on day 1. Solriamfetol significantly affected the frontal EEG delta power during arena habituation (TreatmentXTime interaction effect F(24, 191)=2.422, \*\*\**P*<0.001; Treatment effect: ns; Time effect: ns; mixed-effects analysis followed by Šídák’s multiple comparisons test, Frontal: n=5 for Saline and n=5 for Solriamfetol 3 mg/kg, Parietal: n=4 for Saline and n=6 for Solriamfetol 3 mg/kg). **C-D**. Frontal (C) and parietal (D) beta2 power before, during and after the first 5-min habituation to the test arena. Solriamfetol significantly affected the parietal EEG beta2 power during arena habituation (TreatmentXTime interaction effect F(24, 191)=1.688, \**P*<0.05; Treatment effect: F(1, 8) = 0.045, ns; Time effect: F(2.571, 20.46) = 13.51, \*\*\*\**P*<0.0001; mixed-effects analysis followed by Šídák’s multiple comparisons test, Frontal: n=5 for Saline and n=5 for Solriamfetol 3 mg/kg, Parietal: n=4 for Saline and n=6 for Solriamfetol 3 mg/kg). **E-F**. Frontal (E) and parietal (F) beta2 power during the 10-min NOR acquisition phase on day 2. Solriamfetol significantly affected the parietal EEG beta2 power during the learning phase (TreatmentXTime interaction effect F(29, 226) = 1.634, \**P*<0.05, Treatment effect: F(1,8) = 0.980, ns; Time effect: F(2.297, 17.90) = 15.51, \*\*\*\**P*<0.0001; mixed-effects analysis followed by Šídák’s multiple comparisons test, Frontal: n=5 for Saline and n=5 for Solriamfetol 3 mg/kg, Parietal: n=4 for Saline and n=6 for Solriamfetol 3 mg/kg). **G-H**. Frontal (G) and parietal (H) fast-gamma power before, during and after the NOR testing phase on day 3. Solriamfetol significantly affected the parietal EEG fast-gamma power during the NOR test (TreatmentXTime interaction effect F(29, 254) = 1.538, \**P*<0.05; Treatment effect: F(1, 9) = 1.581, ns; Time effect: F(3.067, 26.86) = 25.63, \*\*\*\**P*<0.0001; mixed-effects analysis followed by Šídák’s multiple comparisons test, Frontal: n=5 for Saline and n=5 for Solriamfetol 3 mg/kg, Parietal: n=5 for Saline and n=6 for Solriamfetol 3 mg/kg). Data are expressed as mean ± SEM.

On day 1 during the 1^st^ 5 min-habituation phase, i.e., during first exposure to the novel context, Solriamfetol affected the frontal EEG delta power, with a significant DrugXTime interaction: while under saline, EEG delta increased, suggesting gradually increased sleepiness, EEG delta remained stable, or decreased, under Solriamfetol 3 mg/kg (Figure 4A).

Additionally, a DrugXTime interaction effect was also observed on the parietal EEG beta2 power, with higher beat2 activity observed in Solriamfetol 3 mg/kg group in early habituation phase (Figure 4D). EEG beta2 oscillations are reported to be associated with novelty recognition^24^.

On day 2 during the acquisition phase (10 min exposure to two identical objects in the habituated context), the parietal EEG beta2 power showed a transient increase in the Saline group above Solriamfetol 3 mg/kg values, but then realigned with values of the Solriamfetol 3 mg/kg group (Figure 4F).

Strikingly, on day 3 when exposed to the novel object during the 10 min NOR test, Solriamfetol 3 mg/kg-treated mice showed increased parietal EEG fast-gamma power values relative to Saline-treated mice (Figure 4H), suggesting higher levels of concentration. Together these data suggest that Solriamfetol 3 mg/kg decreases EEG markers of sleepiness, but enhanced EEG indices of alertness and novelty discrimination in novel contexts, and concentration under cognitive challenge.

### Solriamfetol leads to sustained EEG markers of alertness and attention following the NOR test

After the end of the NOR test, EEG delta and fast-gamma powers were assessed for a total of 30 min. Solriamfetol 3 mg/kg-treated mice showed no difference in average delta power across the full 30 min interval (Figure 5A-B) but exhibited significantly higher frontal fast-gamma activity (Figure 5E). Furthermore, when the 30 min interval was divided into 10 min time-bins, frontal delta power showed a significant TreatmentXTime interaction effect, with the frontal delta power curve gradually increasing under saline, while it decreased under Solriamfetol 3 mg/kg, suggesting that the drug was able to decrease post-test sleepiness (Figure 5C). Moreover, both parietal and frontal EEG fast-gamma powers showed a significant Solriamfetol 3 mg/kg effect, suggesting that the drug conferred a positive and sustained effect on concentration and cognitive activity (Figure 5G-H).

**Figure 5.**
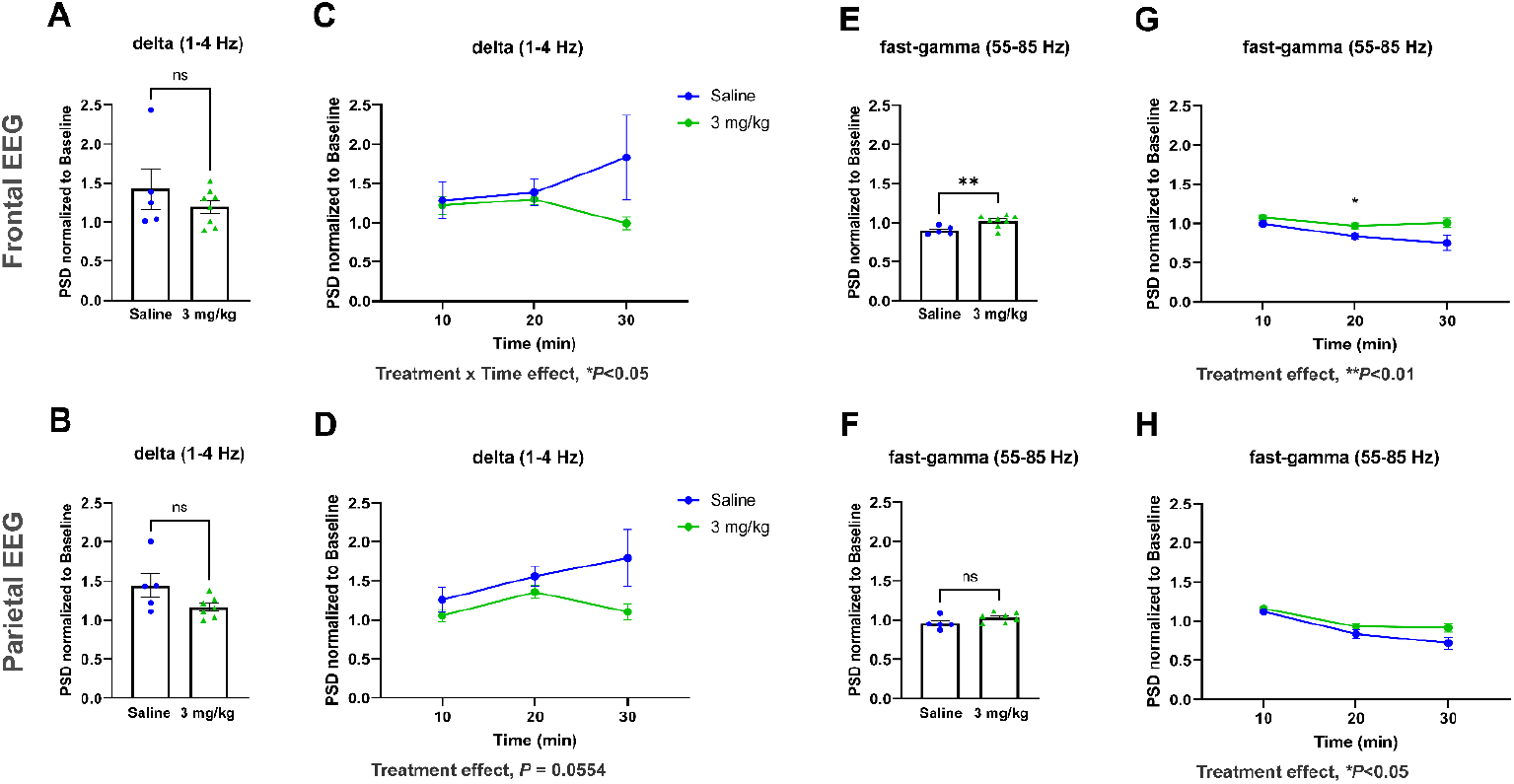
Effects of Solriamfetol (3 mg/kg) on the waking EEG of C57BL/6J mice in the 30 min upon return to the home cage following the NOR test. A-B. Frontal (A) and parietal (B) delta power during the full 30 min following the NOR test. (Two-tailed, unpaired t-test, *P*>0.05). **C-D**. Frontal (C) and parietal (D) delta power following the NOR test analyzed in 10 min time bins. The frontal delta EEG showed a significant TreatmentXTime interaction effect (F(2, 21) = 3.980, \**P*<0.05; treatment effect: F(1, 11) = 1.893, *P*>0.05; time effect: F(1.148, 12.06) = 0.546, *P*>0.05; mixed-effects analysis followed by Šídák’s multiple comparisons test). The parietal delta EEG power only showed a trend toward a teatment effect (TreatmentXTime interaction effect), F(2,19) = 2,107, *P*>0.05; Treatment effect: F(1, 10) = 4.699, *P*=0.0554; Time effect: F(2, 19) = 2.107, *P*>0.05; mixed-effects analysis followed by Šídák’s multiple comparisons test, A-D: Frontal: n=5 for Saline and n=8 for Solriamfetol 3 mg/kg, Parietal: n=5 for Saline and n=7 for Solriamfetol 3 mg/kg). **E-F**. Frontal (E) and parietal (F) fast-gamma powers during the full 30 min directly following the NOR test. Solriamfetol significantly increased frontal fast-gamma activity (\*\**P*<0.01 vs Saline group, two-tailed, unpaired t-test). **G-H**. A significant effect of Solriamfetol 3 mg/kg vs saline on frontal (G) and parietal (H) fast-gamma power was also seen following NOR testing when the 30 min were analyzed by 10 min time bins (Frontal fast-gamma EEG power: Treatment effect: F(1, 11) = 10.13, \*\**P*<0.01; TreatmentXTime effect F(2, 21) = 1.785, *P*>0.05, Time effect: F(1.331, 13.98) = 6.205, \**P*<0.05. Post-hoc test: \**P*<0.05; Solriamfetol 3mg/kg vs Saline group; Parietal fast-gamma EEG: Treatment effect: F(1, 10) = 5.358, \**P*<0.05; TreatmentXTime interaction effect F(2, 19) = 1.547, *P*>0.05; Time effect: F(1.792, 17.02) = 30.09, \*\*\*\**P*<0.0001; mixed-effects analysis followed by Šídák’s multiple comparisons test, E-H: Frontal: n=5 for Saline and n=8 for Solriamfetol 3 mg/kg, Parietal: n=5 for Saline and n=7 for Solriamfetol 3 mg/kg). Data are expressed as mean ± SEM.

## Discussion

Solriamfetol is a relatively new CNS stimulant, only used since 2019 and exclusively for EDS in narcolepsy and OSA. Solriamfetol has several advantages over other stimulants, notably that it is not associated with psychomotor effects, does not induce behavioral stereotypies and anxiety-related behaviors in mice, as observed for instance with amphetamines and modafinil^25^.

Solriamfetol mode of action is thought to be primarily dopaminergic and noradrenergic acting as a DNRI (Baladi et al., 2018). More recently, Solriamfetol was reported to additionally act *in vitro* as a TAAR1 agonist, unlike modafinil and bupropion, but in common with amphetamine and methamphetamine, and as a 5HT1A agonist^13,15^. TAAR1 partial agonists were shown to dose-dependently increase wakefulness in rats and mice, and reduce NREM and REM sleep, suggesting a role for TAAR1 in the control of vigilance states and cortical activity^26,27,28^.

Our initial study demonstrating powerful dose-dependent wake-promotion by Solriamfetol, showed that at 150 mg/kg the drug has an initial surprising effect, inducing for the first 30–40 min a waking state during which mice are alert and responsive, with open eyes, but without significant spontaneous locomotor activity. This contrasts with other CNS stimulants, such as amphetamines, which induce hyperactivity and behavioral stereotypies. These effects were also seen at lower doses, including 50 mg/kg, but were not seen at 3 mg/kg.

Here we show that at 3 mg/kg Solriamfetol is able to enhance performance in short (OBAT) or long-term (24 h, NOR test) memory tests, as well as to enhance spatial memory in the Morris Water maze test. In all three tests, we observed that Solriamfetol cognitive enhancing effects are most apparent in the late phase of the test (when the probe times of 10, 5 or 1 min, for NOR, OBAT and MWM tests respectively, are dissected into shorter time-bins). These results suggest that Solriamfetol enhances cognitive performance by prolonging and maintaining attention. Our EEG data support this interpretation as reflected in a decrease in delta power and increase in fast-gamma power during the probe test of the NOR task, when distinction between the novel and the familiar object takes place. Upscaling of EEG fast-gamma power is known to be associated with cognition in both animals and humans^29 30^.

During exposure to the novel arena on the 1^st^ day of the NOR test (habituation phase), we also observed enhanced parietal beta2 frequencies after administration of Solriamfetol 3 mg/kg in comparison to saline. This increase was most pronounced during the first min of the exposure to the novel context. Interestingly, hippocampal beta2 oscillations have been described as important novelty-detection feature ^31^. Similar to our findings, beta2 enhancement was described to be most prominent during the first two min after novelty exposure^24^. In our earlier study, in which we exposed mice to Solriamfetol 150 mg/kg, a marked increase in beta2 power was also observed, starting about 1 h after injection and lasting for 2 h, and was a Solriamfetol-specific effect, as not seen after d-amphetamine or modafinil administration^11^.

Another observation was that parietal beta2 power decreased across time during the 1^st^ habituation phase, as consistent for a novelty-detection feature. In contrast, during the NOR test phase exploration, as the animal learns to discriminate the novel from the familiar object, parietal fast-gamma activity remains relatively stable throughout the 10 min NOR test, and significantly more powerful under Solriamfetol, in consistency with the observed behavior. Indeed, when mice are exposed to a novel context, novelty reduces with time, and so does parietal beta2 power. In contrast, during the test, mice explore both familiar and novel objects, a task requiring sustained attention to achieve accurate discrimination between two relatively similar objects.

Our study supports previous reports on the cognitive-enhancing effects of psychostimulants on normal subjects. The first significant demonstration that the cognitive-enhancing effects of psychostimulants are not exclusive to patients with hyperactivity disorder was published in 1980, with a report that d-amphetamine, a stimulant used for the treatment of ADHD and narcolepsy, not only alleviated ADHD symptoms in ADHD-afflicted male children, but also enhanced vigilance and learning in normal male children, as well as in normal male adults^32^.

Studies on modafinil showed an enhancement of working memory and short-term memory in mice^33-35^. However, unlike Solriamfetol, modafinil did not have an effect on long-term memory, assessed by the MWM test^35^.

Another psychostimulant reported to have cognitive-enhancing properties is methylphenidate, used in the treatment of ADHD. It is interesting to note that the effect of methylphenidate is contingent upon its dose, brain region, and neuronal population. Like Solriamfetol, a low dose of methylphenidate improved working memory. However, a high dose impaired it^36^.

In mice, fear memory was improved at low doses but impaired at high doses of d-amphetamine, cocaine, or modafinil^37,38,39,40.^ The behavioral-activating effects associated with elevated doses of psychostimulants are absent from low doses of stimulants. This may be explained by a modest increase in NA/DA efflux in the nucleus accumbens and medial septum at lower doses. This, in turn, could reduce the liability for drug abuse^41-43^. Our findings, along with the aforementioned studies, provide evidence that the dose is a crucial factor in determining the cognitive effects of Solriamfetol and other psychostimulants.

Like most psychostimulants, a clinical study suggested that Solriamfetol may prove to be an efficacious treatment for ADHD in adults^44^. We believe that Solriamfetol may have an inverted U-shaped response, where the effective and optimal doses for the cognitive enhancing effect is much lower than the psychomotor effect. And this effect may be mainly mediated by the ability of Solriamfetol to bind to dopamine transporter and increase DA levels in specific brain areas such as the prefrontal cortex while not affecting the DA levels in the striatum, a brain area associated with locomotion. Levin et al.^45^ described an inverted-U shaped model for the relationship between the dopamine levels in the prefrontal cortex and cognitive performance.

In this study, we were able to provide evidence that Solriamfetol at the low dose of 3 mg/kg enhances alertness, novelty-discrimination and cognitive performance in WT mice. A dose of 3 mg/kg in mice is equivalent to ∼0.243 mg/kg in humans according to Nair et al^46^, corresponding to ∼15-20 mg in a subject of 60-70 kg. As the cognitive-enhancing effect may last for 1-3 h, healthy humans might need 2-3 doses of 15 mg a day, or an effective daily dose of 30-45-60 mg, to enhance performance throughout the day. In comparison, narcolepsy patients start with 37.5 mg and increase to 75 mg, with a maximal effective dose against EDS of 150 mg (75 mg twice daily). Therefore, Solriamfetol dosage to increase cognitive performance in healthy subjects may be 3-5-fold lower than dosage prescribed for EDS in narcolepsy and OSA.

Our data may contribute to the improvement and adjustment of the use of Solriamfetol and other psychostimulants in narcolepsy and other sleep disorders. They support the notion that, like other stimulants, the therapeutic effect of Solriamfetol is highly dose-dependent, and lower doses are beneficial for cognitive performance in mice that are not cognitively impaired. The mechanism of action may be attributable to its capacity to inhibit dopamine transporter. The exact mechanisms of action in the human body remain however incompletely solved and warrant further investigations. It will additionally be of great importance to determine the brain regions implicated in Solriamfetol action.

## Data Availability Statement

Correspondence and data requests should be addressed to Anne Vassalli.

## Acknowledgment

We would like to thank Richie Kalusivikako for assistance during behavioral experiments.

## Author Contributions

MH performed all experiments; MT and AV designed research; MH and LYC contributed new analytic tools; ST contributed behavioral experimental design; MH, ST, LYC, AV and MT analyzed data; and MH, AV, and MT wrote the paper.

